# Deterministic Critical Community Size for the SIR System and Viral Strain Selection

**DOI:** 10.1101/2020.05.08.084673

**Authors:** Marcílio Ferreira Dos Santos, César Castilho

**Affiliations:** Núcleo de Formação de Docentes, Universidade Federal de Pernambuco, Caruaru, PE CEP 55014-900 Brazil; Departamento de Matemática, Universidade Federal de Pernambuco, Recife, PE CEP 50740-540 Brazil

**Keywords:** Epidemics, Virulence, SIR Modes, Evolution Theory

## Abstract

In this paper the concept of Critical Community Size (CCS) for the deterministic SIR model is introduced and its consequences for the disease dynamics are stressed. The disease can fade out after an outburst. Also the principle of competitive exclusion holds no longer true. This is exemplified for the dynamics of two competing virus strains. The virus with higher *R*_0_ can be eradicated from the population.

## 1. Introduction

The Critical Community Size (CCS) of a infectious disease is defined as the minimum size of a closed population within which the disease’s pathogen can persist [4, 5]. When the size of the population is smaller then the CCS, the low density of infected hosts causes the extinction of the pathogen after an epidemic outbreak. In this case the disease is said to fade out [3, 2]. Classical SIR deterministic models for direct contact viral diseases [6] fail to capture the fade out phenomena: either the number of infected converges to an endemic equilibrium after successive outbreaks or it disappears without any outbreak; the fate of the disease depending on a bifurcation parameter, called the basic reproductive number *R*_0_ [11, 9].

One of the reasons for the SIR breakdown to capture the disease fade out for small populations is the use of real numbers to count individuals. While being a good approximation for large populations, counting individuals using real numbers has dramatic consequences when the number of individuals within a particular compartmental class becomes smaller then one and therefore extinct: the SIR model fails to capture the small population extinction. In this paper extinction is incorporated into the model. When one of the variables representing the compartmental classes becomes smaller then one it is immediately set to zero. This has two important consequences. First the concept of Critical Community Size appears naturally. Second the principle of competitive exclusion no longer holds.

In a realistic context, when the number of individuals is too small, the probability of disease eradication is high. This Allé The paper is organized as follows. In SECTION 2 the CCS for the classical SIR model with constant population is defined. The definition follows directly: for a population of *N* individuals the disease will be eradicated if the density of infected individuals becomes smaller then 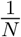. This simple fact, that can be also interpreted as a consequence of the Alle effect [1, 8], allows for the determination of the minimum viable population [12] for a disease. This minimum viable population does not dependent on the disease *R*_0_ value, instead it depends non trivially on the parameters reflecting the scaling properties of the SIR system. Curves for the CCS in terms of the parameters of the SIR system are exhibited.

In SECTION 3 we study the consequences of the fade out on the dynamics of two different competing virus strains. It is numerically shown that the Principle of Competitive Exclusion (PCE) [7] is no longer true: A strain with smaller *R*_0_ can eliminate one with higher *R*_0_. On section 4 we draw our conclusions

## 2. SIR model and Critical Community Size

Let *S*(*t*), *I*(*t*) and *R*(*t*) denote the number of susceptibles, infected and removed at time *t* respectively and *N*(*t*) = *S*(*t*) + *I*(*t*) + *R*(*t*) be the total number of individuals. The SIR model states that

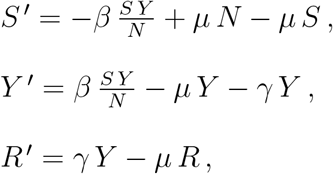

where *β* is the infection rate, *γ* is the clearance rate and *μ* is the mortality rate (assumed equal to the birth rate). Adding the equations it follows that *N*(*t*) is a first integral. Introducing the densities 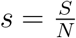, 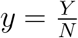 and 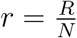, equations become

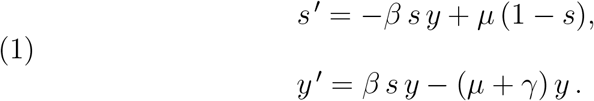

The equation for *r*(*t*) is omitted since *r*(*t*) = 1 – *s*(*t*) – *y*(*t*).

The qualitative dynamics of the above system is determined by the bifurcation parameter

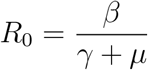

called the disease basic reproductive number. It represents the average number of cases caused by one infected individual in a totally susceptible population. The following are well known facts [6].

i. If *R*_0_ < 1 the dynamical system has only one equilibrium point *E*_0_ = (1,0) called the disease free equilibrium point. *E*_0_ is a globally stable critical point.
ii. If *R*_0_ > 1 the dynamical system has two equilibria. *E*_0_ an unstable critical point and 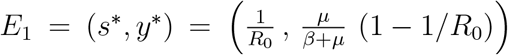 a globally stable critical point. The global stability of *E*_1_ can be proved using the Lyapounov function [10]

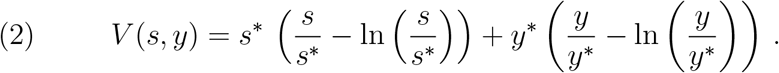

Since *N* represents the total number of individuals one must have 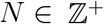. Accordingly, the smallest possible value for the densities *s*(*t*), *i*(*t*) and *r*(*t*) is 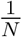 and if any of the densities becomes smaller than 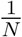 the correspondent population becomes extinct. This simple fact is not usually taken into consideration and it is of utmost importance for what follows.

*Remark* 2.1. Introducing a new independent parameter defined as *τ*(*t*) = *μ t* the SIR system becomes

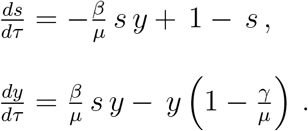

Therefore, the qualitative dynamics can be studied by fixing a *μ* value and considering the *γ* and *β* parameters as measured in *μ*-units.

### 2.1. Fade out for the SIR model

The disease is said to fade out if it disappears in a finite time after some epidemic outbreak. For the deterministic SIR model this is only possible if for some time instant the number of infected individuals is smaller then one, or equivalently if *y*(*t*) < 1/*N*.

#### Definition 2.2.

Let *R*_0_ > 1. The disease will *fade out* if for some time instant *t*\*

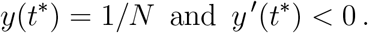

The *R*_0_ > 1 condition allows for the possibility of epidemic outbreaks.

In what follows the fade out phenomena for the SIR will be characterized for the initial conditions

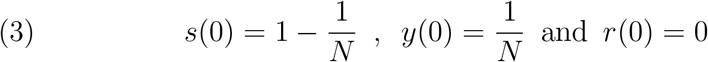

representing the invasion of a totally susceptible community by only one infected individual. The characterization of the fade out for the SIR model is done by noticing that the local minima for the *y*(*t*) curve form a monotonic increasing sequence (see Appendix A for a proof of this fact). Therefore, in the case it exists, the first local minimum of *y*(*t*) is also the global minimum for *t* > 0.

#### Definition 2.3.

Assume 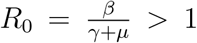. Define Ψ = Ψ(*β*, *γ*, *μ*, *N*) as the value of the first minimum of *y*(*t*) with *t* > 0, for the initial conditions 3. If *y*(*t*) has no minimum value for *t* > 0 set 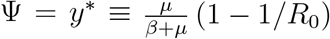.

#### Definition 2.4.

Let *γ*, *β*, *μ* all fixed and positives. 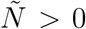 is called the *Deterministic Critical Community Size* if

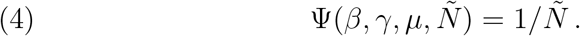

If the population is smaller then 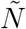 the disease will fade out and if the population is greater or equal to 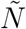 the number of infected individuals will converge to its limit value *y*\*. The numerical procedure for the determination of the CCS is straightforward: Let *α*, *β* and *γ* such that *R*_0_ > 1. Fixing a value for *N*, integrate system (1) up to the first *y*(*t*) local minimum (with *t* > 0). If the minimum value is equal to 1/*N* then *N* is the CCS. If not, change the *N* value and proceed on the same way. A bisection method can be used to determine the CCS.

The CCS dependence on the parameters *γ* and *β* is showed on figure

The numerical results are exhibited in figure (1) for *μ* =1.0 × 10^−5^. As an example notice that considering a population of 10000 persons all the parameters values in the yellow area will drive the disease to extinction in figure (3). The disease will persist for all parameters values outside the yellow region. Also notice that the level curves coalesce very near the *R*_0_ = 1 curve for larger values of the clearance rate. The consequence of this coalescence is that the CCS values for diseases with *R*_0_ ≈ 1 can assume any value. The numerical results shows that the *R*_0_ value, while determining the global dynamics of the SIR system, does not determine the CCS.

**Figure 1.**
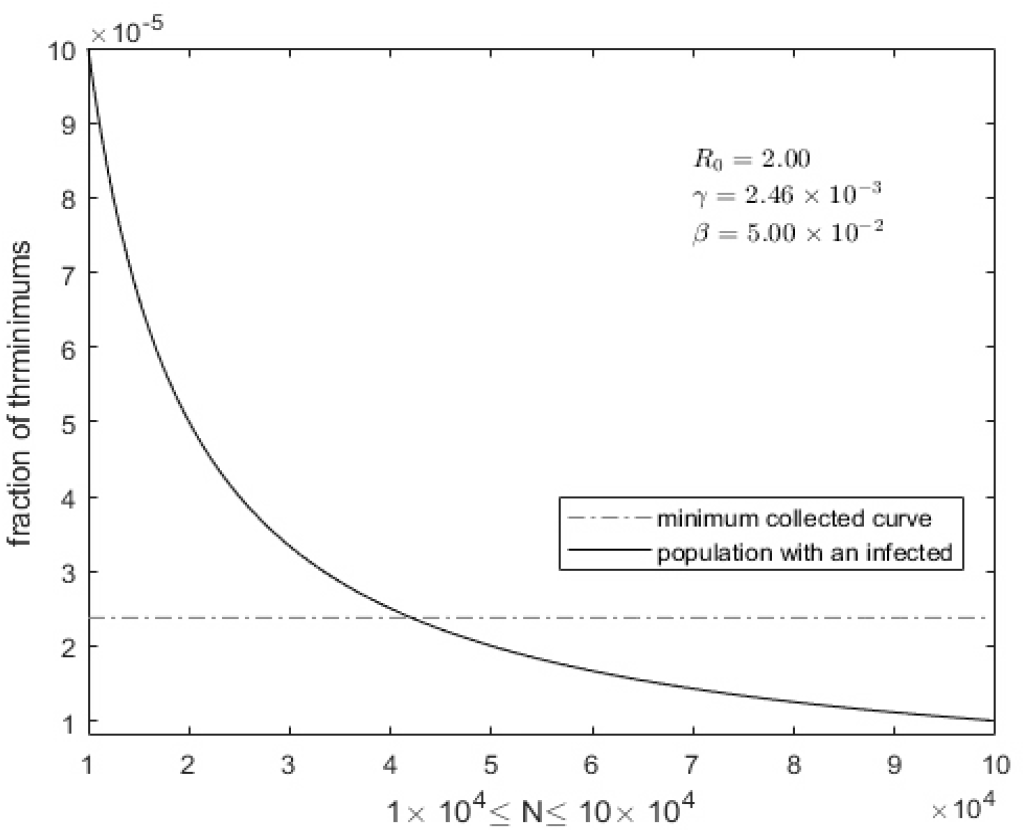
Numerical determination of the Critical Community Size (CCS).

**Figure 2.**
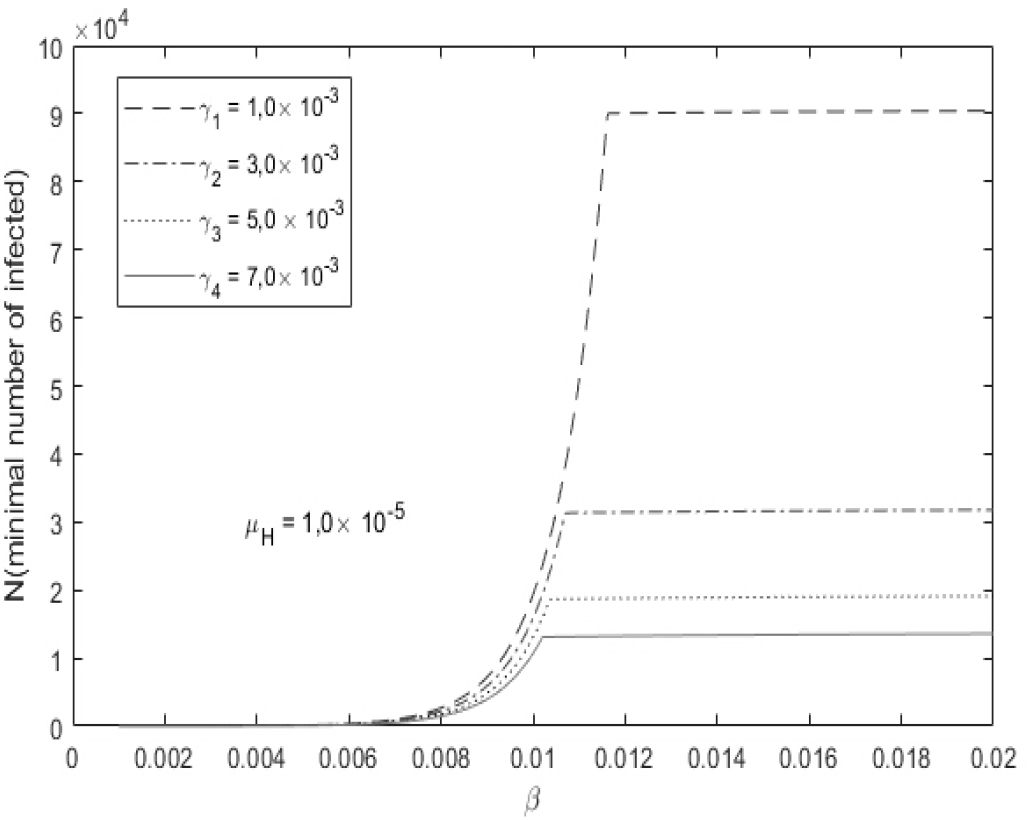
CCS as a *β* function. *γ* values are indicated on figure.

**Figure 3.**
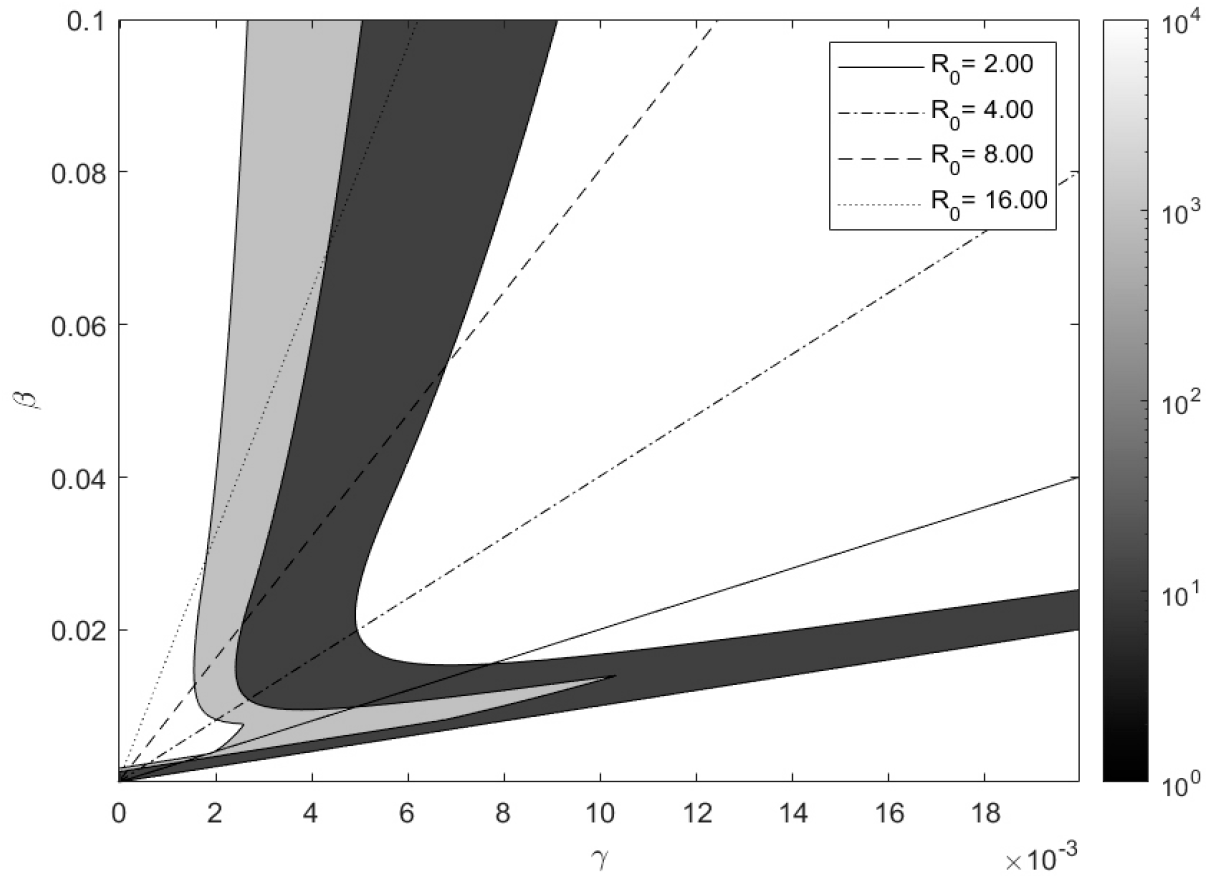
Critical Community Size (CCS) as a function of *γ* and *β*. The level curves represent the CCS values. The mortality rate is *μ* = 1.0 × 10^−5^.

## 3. Viral Competition under Environmental Selective Pressure

In this section the implications of the CCS for a system with two competing virus strains will be explored. The main fact here is that the Principle of Competitive Exclusion (PCE) will no longer holds.

Let *S*(*t*), *I_i_*(*t*) (*i* = 1, 2), *R*(*t*) denote the numbers of individuals of susceptibles, infected by strain *i* = 1, 2, and removed at time *t* respectively. Let *N*(*t*) = *S*(*t*) + *I*_1_(*t*) + *I*_2_(*t*) + *R*(*t*) be the total number of individuals on the population at time *t*. Let

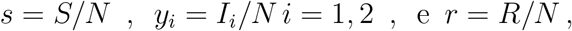

be the respective densities. 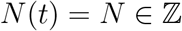 since it represents the total number of individuals on the population. Accordingly, the smallest possible value for the densities *s*(*t*), *i_i_*(*t*) and *r*(*t*) is 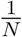. In order to make the discussion more general, an Allee effect will be introduced: If *I_i_*(*t*) ≤ *ρ* for some critical community size 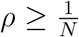 the disease will be considered extinct. *ρ* is expected to depend on the population density, mixing, etc.

The two-strain model is

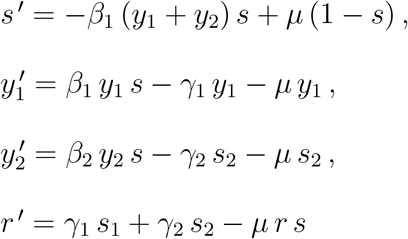

The CCS for the above system is difficult to determine since the minima are no longer monotonically ordered and the parameter space is larger. To show that the basic reproductive number of the virus is no longer the unique factor deciding the outcome of the viral competition the first minima of *I*_1_(*t*) and *I*_2_(*t*) are calculated for the following situation: The parameters of the virus-1 are held constant at *R*_1_ = 3, 20, *β*_1_ = 9, 0 × 10^−3^, *μ*_1_ = 5 × 10^−5^. For the virus-2 *R*_2_ = 3, 20, *μ*_2_ = 5 × 10^−5^ and *β*_2_ varying from 4,0 × 10^−3^ to 19, 0 × 10^−3^. The noticeable point here is the inversion of the infected minima curves. For *β*_2_ < 3.0 × 10^−3^ the minima of *y*_1_ are greater then *y*_2_ and for *β*_2_ > 3.0 × 10^−3^ the opposite holds. For a population threshold of 1 × 10^−5^ the virus-2 is eliminated if *β*_2_ < 3.0 × 10^−3^ and the virus-1 can be eliminated if *β*_2_ > 3.0 × 10^−3^. Notice the value o *R*_0_ has been held constant for both viruses. The simulation is not contradicting the PCE since both viruses have the same *R*_0_ value.

*Remark* 3.1. The example in figure (5) began with two people with a virus-1 and virus-2 strains each. Although after some time the strain with *β*_1_ = 12 × 10^−3^ have no one infected individual. Therefore we can conclude that one strain go to extinction.

**Figure 4.**
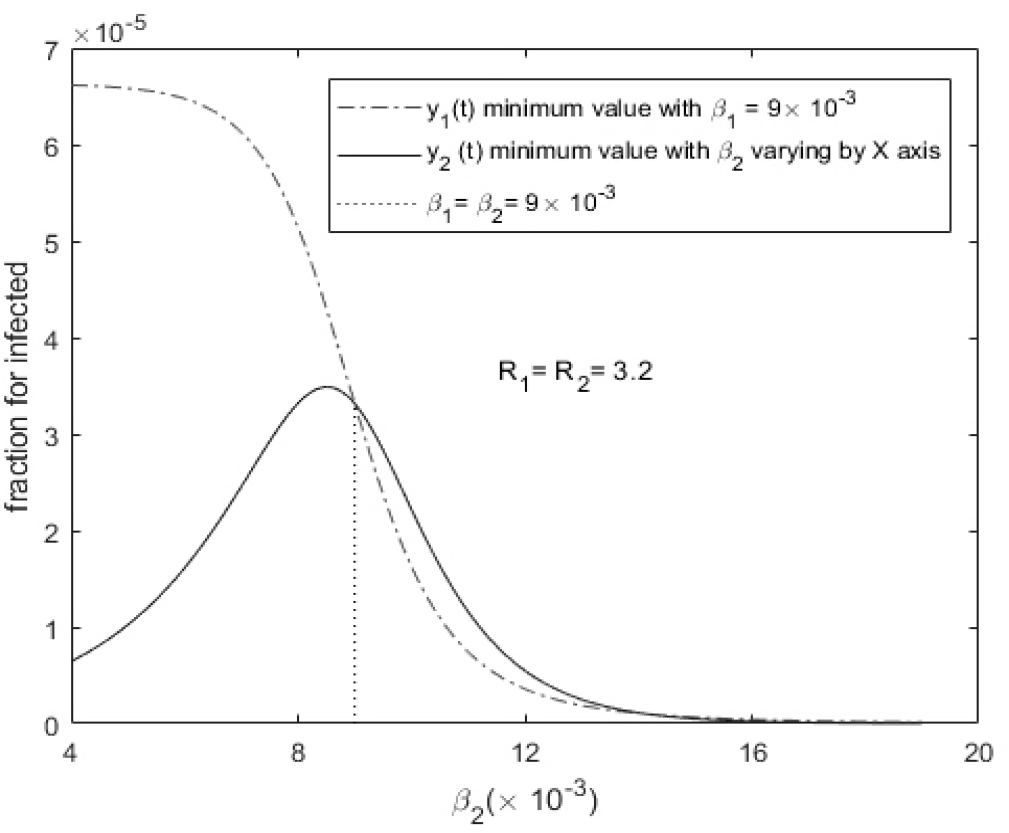
First minima for the two virus strains as a *β*_2_ function and fixed *μ_H_* = 5 × 10^−5^.

**Figure 5.**
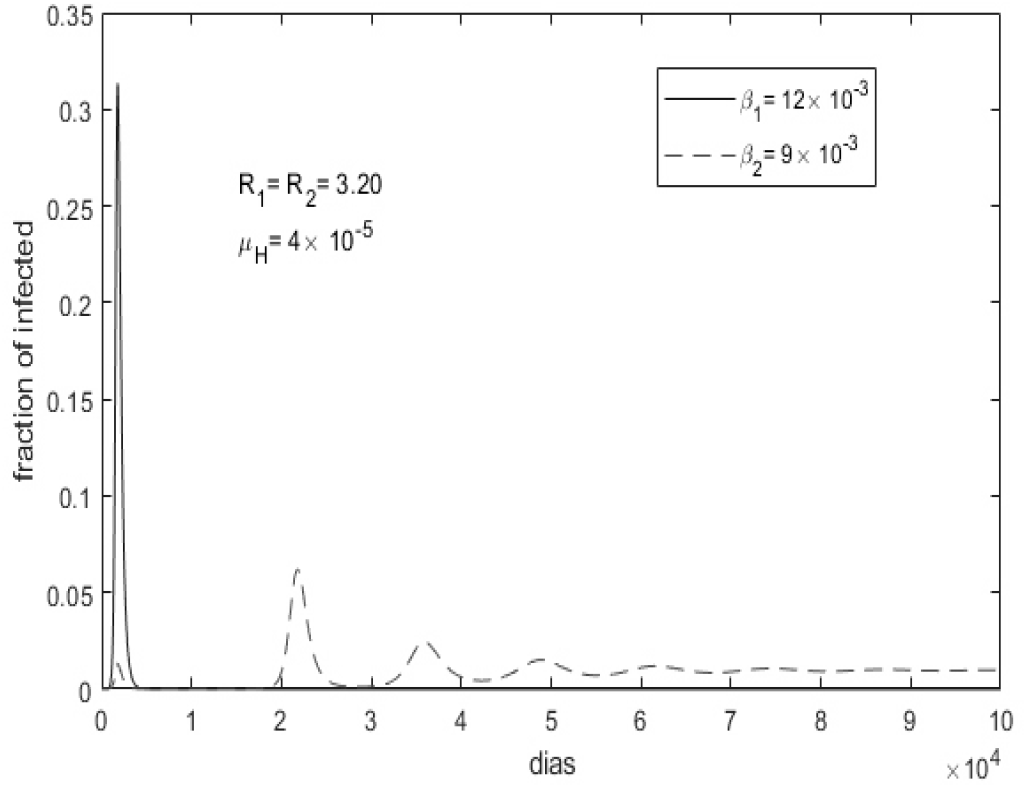
Elimination of the virus-1 after passing trough its first minimum.

**Figure 6.**
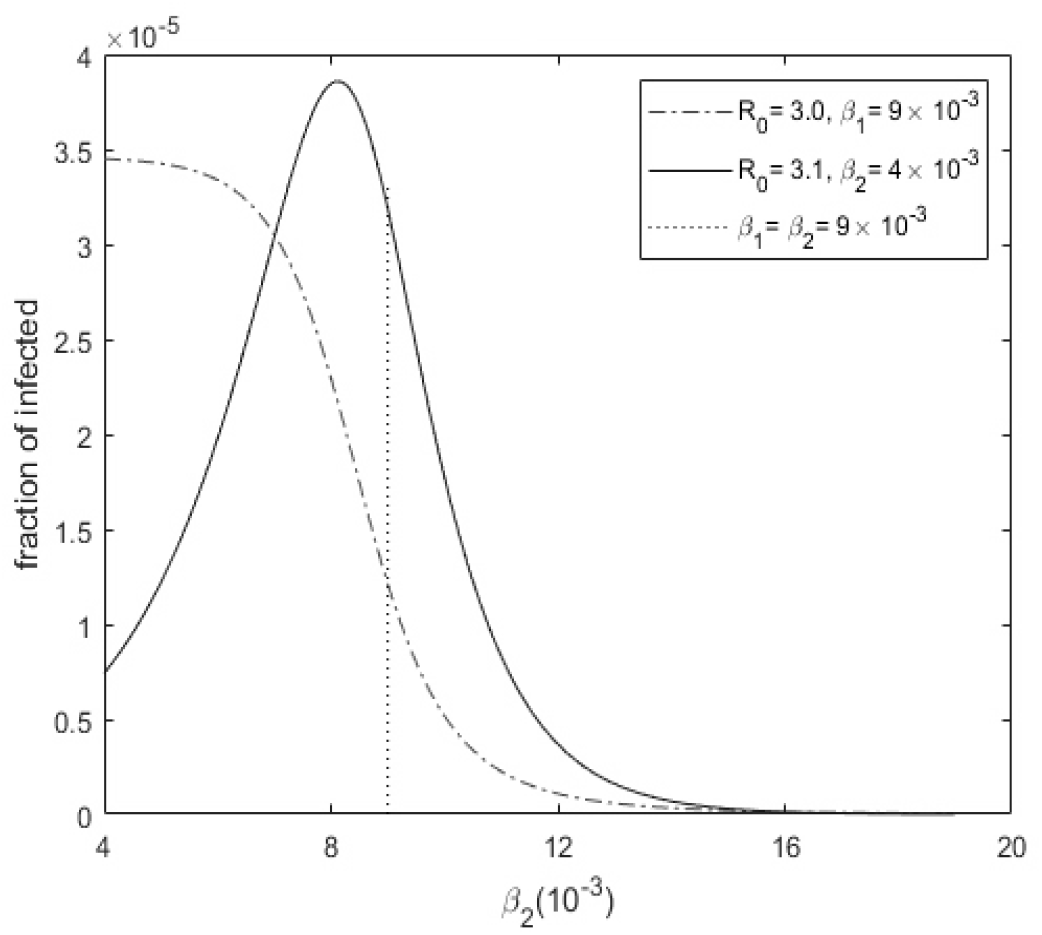
First minima for the two virus strains as a *β*_2_ function.

**Figure 7.**
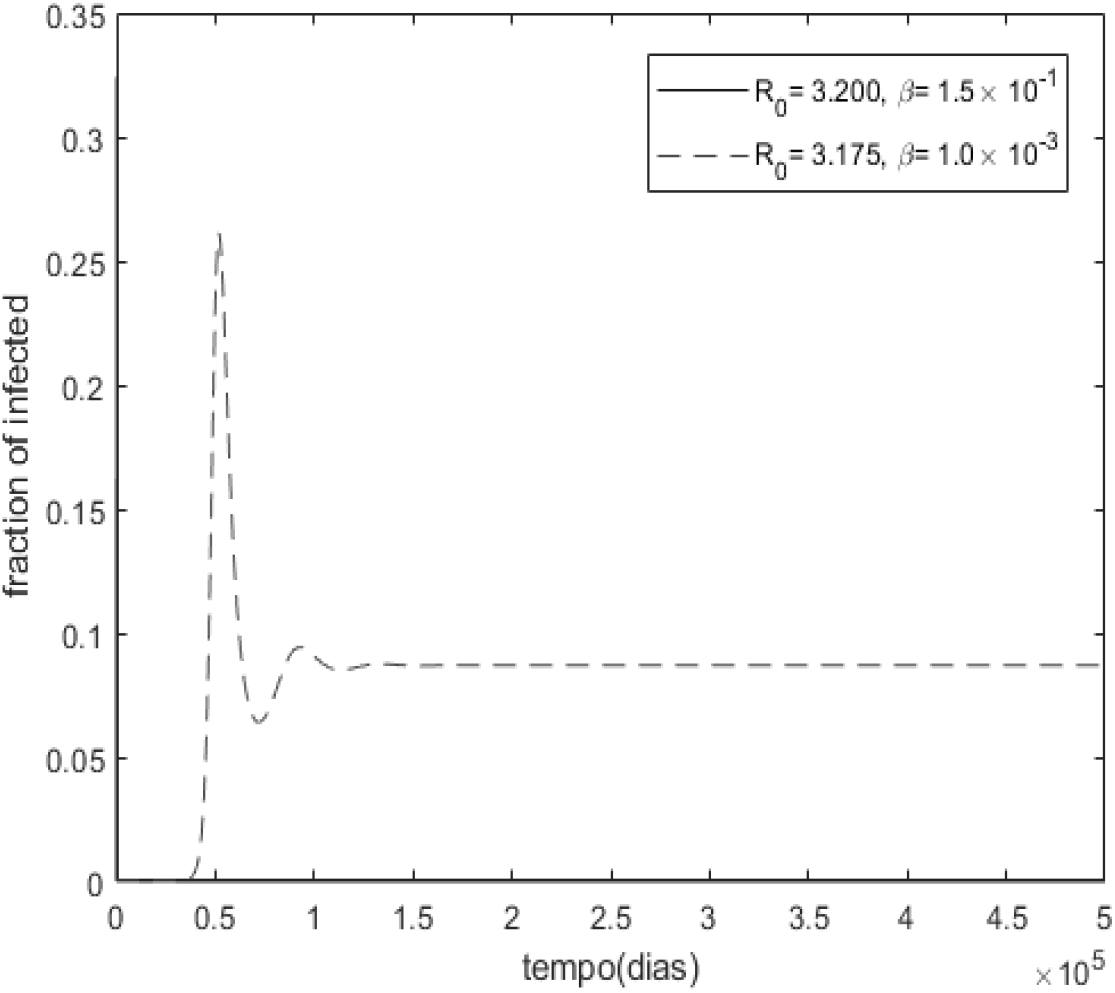
Elimination of the virus-1 after passing trough its first minimum.

The next simulation shows that the value of the first minimum can be smaller for the virus with greater *R*_0_ leading to its eradication and the permanence of virus with smaller *R*_0_ in contradiction to the PCE.

## 4. Conclusions

The concept of a deterministic Critical Size Community, as introduced in this paper, provides an alternative potential mechanism for the disease fade out phenomena. It also allows the characterization of the disease persistence within a community in terms of population size and disease parameters. This characterization is shown in Figure (). The graphic shows for example, that two diseases with the same basic reproductive number *R*_0_ can have rather different dynamics. This is, from a mathematical point of view, consequence of the non-linear scaling properties of the SIR system and has drastic consequences for the disease persistence.

The results also provide evidence that the Critical Community Size is an important component for the viral competition and natural selection processes. While for the PCE, the virus with higher *R*_0_ will eliminate the other viruses with smaller *R*_0_, the concept of CCS shows that the dynamic is in fact more complex. The CCS allows for the extinction of any virus population reaching the minimum threshold value. While still predicting that only one virus type will persist in the long term, it shows that the survivor type needs no longer to be the one with the highest *R*_0_.

## 5. APPENDIX A

### Proposition 5.1.

*The time ordered local minima of y*(*t*) *form a monotonic increasing sequence. Analogously the time ordered local maxima form a monotonic decreasing function*.

*Proof*. Since 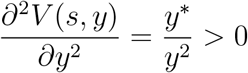 the *E*_1_ Lyapounov function is convex in *y* [10].

Let *y*(*t*_1_) and *y*(*t*_2_) be two consecutive local minima of *y*(*t*). Since they are minima *y*′(*t_i_*) = 0 and *y*″(*t_i_*) > 0, *i* = 1, 2. This implies 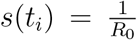 and *y*(*t_i_*) < *y*\*, *i* = 1,2. Since *V*(*s*, *t*) is a Lyapounov function, it is a decreasing function along the flow. Therefore *t*_2_ > *t*_1_ implies that *V*(1/*R*_0_, *y*(*t*_2_)) < *V*(1/*R*_0_, *y*(*t*_1_)) Also, *E*_1_ is a global minimum of *V*(*s*, *y*), it follows by convexity that *V*(1/*R*_0_, *y*) is a *y* decreasing function for *y* < *y*\* and therefore *y*(*t*_2_) > *y*(*t*_1_). The proof for the maxima is similar.

